# Prolonged Antibiotic Treatment Generates a Fluoroquinolone Resistant Gut Microbiome and Collateral Multi-Drug Resistance

**DOI:** 10.1101/467001

**Authors:** Vadim Dubinsky, Leah Reshef, Nir Bar, Keren Rabinowitz, Lihi Godny, Hagit Tulchinsky, Uri Gophna, Iris Dotan

## Abstract

The majority of the gut microbiome develops antibiotic resistance via point-mutations in addition to collateral resistance whereas its density is only moderately decreased following long-term antibiotic treatment.

**ABSTRACT:** Antibiotic resistance in bacterial pathogens represents a growing threat to modern medicine. Limitation of lengthy and broad-spectrum antibacterial treatment regimens is generally recommended. Nevertheless, some conditions may require prolonged antibiotic treatment. The effects of such treatments on bacterial communities, specifically their resistome, is yet unknown. Here, we followed a unique cohort of patients with ulcerative colitis who underwent total large bowel resection and the formation of an ileal pouch from their normal small bowel. The majority of these patients tend to develop inflammation of this previously normal small bowel, known as “pouchitis”. Pouchitis is commonly treated with repeated or prolonged courses of broad-spectrum antibiotics. By using metagenomics of faecal samples obtained longitudinally from patients treated with antibiotics for prolonged periods, we hereby show that the majority of their gut commensal bacteria develop antibiotic resistance by point-mutations. In addition, some bacterial species had acquired multidrug resistance loci with genes that confer resistance to the drug used in the treatment (ciprofloxacin) but co-localized with genes encoding extended-spectrum β-lactamases and other resistance-conferring enzymes. We further show that bacterial density in faecal samples is only modestly reduced despite the long-term antibiotic treatment, thereby questioning the current rationale that antibiotic efficacy in treating pouch inflammation is due to the reduction of bacterial load. This study reveals the impact and dynamics of prolonged antibiotic treatment on human gut microbiomes and provides insights that may guide the development of future IBD therapies. It also provides novel insights into bacterial community recovery after cessation of such prolonged treatment, and highlights the increased risk of spreading mobile antibiotic resistance.

## INTRODUCTION

Crohn’s disease and ulcerative colitis (UC) are chronic inflammatory bowel diseases (IBD) with a significant increase in occurrence worldwide [1,2]. As UC is limited to the large bowel, up to 25% of patients with intractable or complicated UC may undergo total large bowel resection with reconstruction of intestinal continuity by creation of a reservoir ("pouch") from the normal small bowel connected to the anus (total proctocolectomy and ileal pouch anal anastomosis, “pouch surgery”) [3]. Up to 60% of these former UC patients may develop inflammation of the previously normal small bowel comprising the pouch (pouchitis) [4]. The etiology of pouchitis [5,6] and of IBD is assumed to involve an aberrant immune response to imbalanced gut microbiota [7] in genetically susceptible individuals [8], following unknown triggering events [9]. The microbiota of patients with a pouch was shown to be less diverse and with changes in abundance of specific taxa (e.g. decrease in *Faecalibacterium* and increase in *Escherichia* genera) compared to that of individuals with an intact colon [10-13].

The first line therapy of pouchitis is antibiotics. A 2-week course of ciprofloxacin, metronidazole or their combination is usually recommended [6,14]. However, a significant proportion of patients may become antibiotic dependent thus requiring repeated courses or prolonged periods of antibiotic therapy [6,14].

Previous studies focusing on the influence of antibiotics on the gut microbiome of healthy adults [15-18] and infants [19,20] demonstrated that even short-term antibiotic use reduces microbiota diversity as well as short and long term stability, while at the same time enriching the microbiome in diverse antibiotic resistance genes (ARG), some of which are encoded on mobile genetic elements. Notably, humans who were not exposed to antibiotics recently, may also harbour ARG [21,22], as well as those that were isolated from Western civilization for thousands of years [23,24] which implies that human microbiome is an ARG reservoir. Previous studies evaluating the effects of antibiotics, focused mostly only on short (mainly 5-7 days) antibiotic treatments [15,16,19,20,25,26] while prolonged antibiotic therapy and its consequences for the human gut microbiome have been rarely assessed.

Here we present a longitudinal study focusing on patients with a pouch who have been treated with antibiotics for prolonged periods of time, including subsets treated for months to years. Clinical follow up and shotgun metagenomic analysis were performed. Antibiotic resistance in the microbiomes of these patients was explored by evaluating point-mutations in target-encoding genes as well as mobile resistance genes to establish the microbial resistance to antibiotics at the community level. Finally, we inferred bacterial density in the fecal samples through metagenomic analysis validated by qPCR. We show that despite long-term antibiotic treatment only modest reduction in bacterial density was observed. Furthermore, although all patients responded well to the antibiotic treatment, they harboured highly resistant gut bacterial communities, with substantial collateral resistance. This work provides a first look into the impact of long-term antibiotic treatment on the human gut microbiome, and also addresses the mechanism of antibiotic therapy in treating a specific form of IBD.

## RESULTS AND DISCUSSION

### Antibiotics and pouch inflammation have a comparable detrimental effect on bacterial diversity

To gain insights into the effects of long-term antibiotic treatment on pouch microbiome, 49 patients after pouch surgery followed longitudinally were assessed (Supplementary Table 1a). We generated and analyzed 234 shotgun metagenomes from fecal samples obtained from these patients (Supplementary Table 1b). Each patient was sampled multiple times (range 2-12 time-points, median 4, Fig. 1a) over the course of 7 months to 5.9 years (median 3.9 years). Most (n=35) patients in our cohort were treated with antibiotics for prolonged periods of time, with a median of 348 days of antibiotic therapy during follow-up (Supplementary Table 1c,d). Samples were obtained either during antibiotic therapy or within one month after an antibiotics course (n=72), abbreviated hereafter as Abx+, or without antibiotic treatment for at least one month (n=162), abbreviated hereafter as Abx- (Fig. 1a, Supplementary Table 1b). We performed taxonomic profiling of the metagenomic reads and measured bacterial diversity and richness at species level. As only patients with pouchitis are treated with antibiotics, while patients with a normal pouch are not, the effects of pouch inflammation and of antibiotic therapy need to be disentangled. For this end we constructed a generalized linear mixed model with several predictors for species richness, including pouch clinical phenotype (physician assessment, see Methods) and antibiotic usage (Fig. 1b). Each individual patient was set as random effect to control for longitudinal sampling. The model indicated that the clinical phenotype of the pouch, time since last antibiotic use, and cumulative antibiotic duration were significant contributors to decreased species richness in pouch microbiomes. In contrast, VSL-probiotics and anti-TNF therapy increased species richness, yet these trends did not reach statistical significance. Calprotectin, a known clinical marker for intestinal inflammation, was not a substantial predictor of bacterial richness. Next we compared the microbiome of patients with acute/recurrent acute, chronic pouchitis and Crohn’s-like disease of the pouch (CLDP) to the microbiome of patients that have never had pouchitis (Fig. 1c, see definition of pouch behavior in Methods), while excluding samples obtained from patients treated with antibiotics during the last six months (n=115 samples, Supplementary Table 1b). As previously reported by us and others [13,27], patients with a history of pouchitis (especially, the chronic pouchitis and CLDP phenotypes) had a lower median diversity (Shannon index of 2.2) compared to patients with a normal pouch (Shannon index of 2.7) (Fig. 1c, Kruskal-Wallis, P<0.05). We then compared bacterial diversity in patients with a pouch that were treated with antibiotics, grouping samples according to the time that elapsed since the last antibiotic dose. As expected, use of antibiotics significantly reduced bacterial diversity: during antibiotic treatment, or within one month post-treatment, a median reduction of 43% in diversity compared to Abx-samples was observed (Fig. 1d, Mann-Whitney, P=2.1×10^-9^, Shannon index of 2.37 and 1.66, respectively), regardless of the pouch phenotype. Importantly, this effect diminished with time after antibiotic discontinuation (Supplementary Fig. 1a, Kruskal-Wallis, P<0.05) and after ≥ 180 days had elapsed since last antibiotic treatment, diversity has recovered to pre-treatment levels. Thus both antibiotics and a history of pouch inflammation have detrimental influences on pouch microbiome diversity. Notably, different antibiotic regimens (ciprofloxacin, metronidazole or a combination of the two) had comparable effects on bacterial diversity (Supplementary Fig. 1b), despite differences in their spectrum of antibacterial activity. Lastly, for antibiotic treated patients there was no discernible cumulative effect of longer periods of antibiotic treatment on bacterial diversity or richness (Spearman r=-0.06 and r=-0.01, P>0.05, Supplementary Fig. 2), which may be expected since previous human studies showed a strong reduction in diversity after very short courses of treatment [15,25,26].

**Fig. 1.**
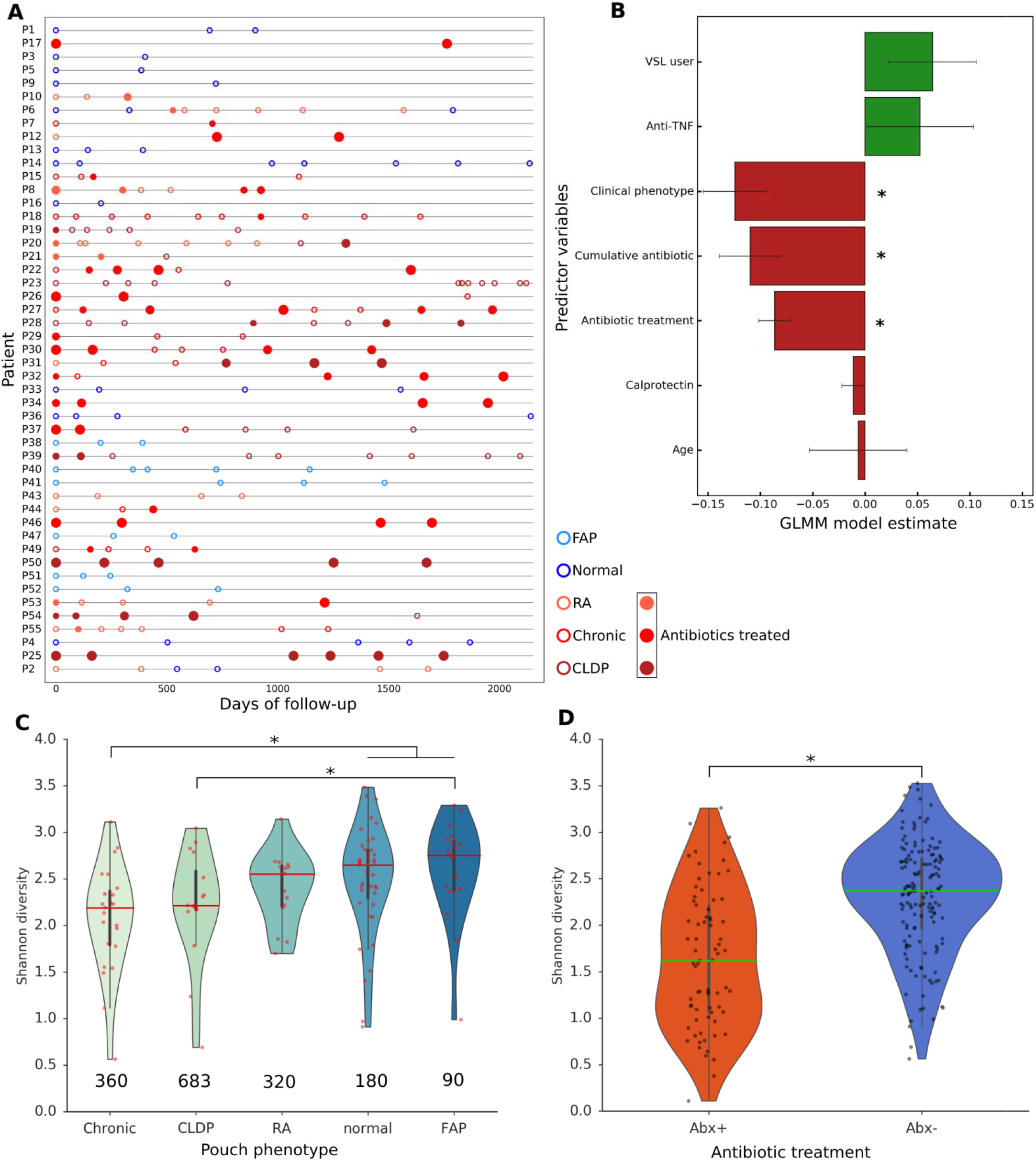
Longitudinal follow-up of pouch patients metagenomes. (a) 231 longitudinal samples from 49 pouch patients were collected over the course of up to six years (x-axis). 73 samples were collected during antibiotic treatment (or up to 1 month after, Abx+); 159 samples were from patients not treated for at least 1 month (Abx-). Colors signify the patients clinical phenotype, open and close circles are Abx- and Abx+ samples, accordingly; Bigger circles correspond to more cumulative days of antibiotic usage, (b) Generalized linear mixed model (GLMM) estimate for species richness based on clinical predictors (y-axis), individual set as random effect; *P < 0.05. Error bars indicate standard error of mean estimates, (c, d) Shannon diversity in the microbiomes based on MetaPhlAn2 taxonomic profiling on the metagenomic reads; samples are separated according to (b) clinical phenotype, samples from patients treated with antibiotics during the last half year were excluded (c) Abx+ and Abx- groups, where all samples were included in this analysis, (c) Statistical comparisons were performed with Dunn’s test to correct for multiple comparisons; Kruskal-Wallis test, *P < 0.05. (d) Mann-Whitney U test, P=2.1e-9. Violin plots whiskers mark observations within 1.5 interquartile range of the upper and lower quartiles. The numbers in right below the plots represent median fecal calprotectin levels for the samples in the corresponding group.

### Stability of the microbiota in patients with pouchitis is low but does not decrease further following long-term antibiotic usage

Stability of the microbiota is defined as the fraction of shared species between two consecutive time points (Jaccard similarity). We hypothesized stability would be relatively low in patients with pouchitis. This may be attributed to the significantly lower volume of the pouch, as well as to previous observations demonstrating that the concentration of bacteria in feces of patients with a pouch is lower compared to the colon of healthy individuals [28]. We found that intra-personal similarity of the microbiota in patients with a pouch is significantly lower than published data for individuals with primary IBD: median stability of 0.32 compared to 0.67 in [32]. However, intra-personal microbial similarity in patients with a pouch was significantly higher than inter-personal similarity (pairwise samples from different patients randomly shuffled, Fig. 2a, Mann-Whitney, P=1.9×10^-20^). This may imply that despite variable time intervals between consecutive samples in our longitudinal dataset and the generally low microbiota stability of patients with a pouch, the concepts of temporal stability and individuality hold true for patients with a pouch. This is in line with previous studies of healthy subjects [29-31] and of patients with IBD [32].

**Fig. 2.**
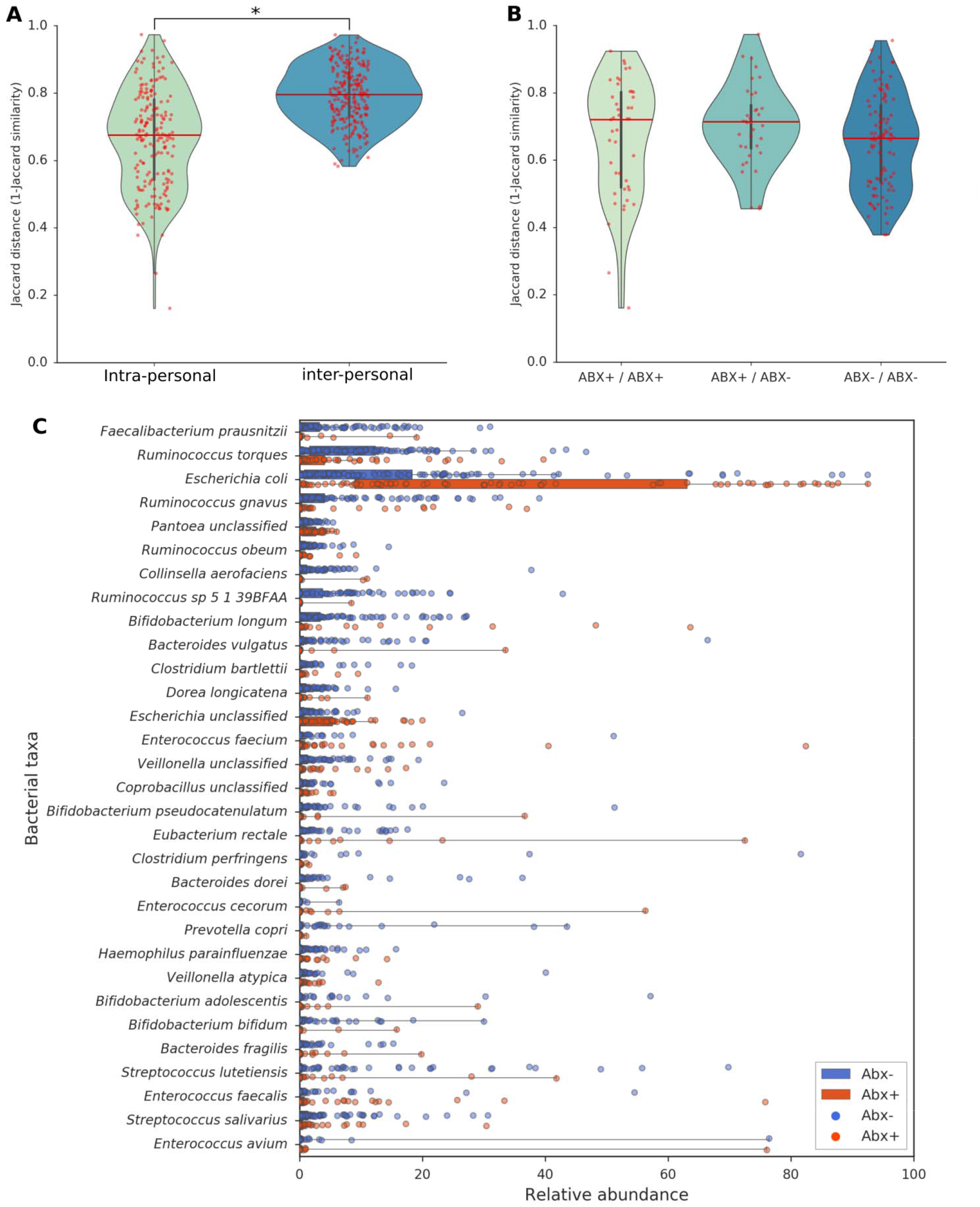
Microbiota stability and differentially abundant bacterial species following antibiotic treatment. (a) Samples are grouped by intra-personal similarity (Jaccard distance of pairwise consecutive samples of each patient) and inter-personal similarity (pairwise samples randomly shuffled), (b) Samples are grouped by: adjacent samples during antibiotic treatment or <30 days after (ABX+ / ABX+), in the absence of antibiotic treatment for >30 days (ABX- / ABX-) and samples whereas one is during treatment and the other in the absence (ABX+ / ABX-). Higher Jaccard distance (1-Jaccard similarity) means lower stability. Mann-Whitney test, *P=7.5e-4, (c) Metagenomic taxonomic profiling compared in Abx+ and Abx- groups. All plotted taxa are significantly different between groups, Mann-Whitney U test, P<0.05 (FDR corrected). Species with a mean relative abundance of < 0.5% across all samples are not shown. Violin plots and boxplot whiskers mark observations within 1.5 interquartile range of the upper and lower quartiles. Red underlines denote taxa that were higher in abundance in Abx-i- group. Full list of species composition in the metagenomes is in Supplementary Table 3

We next sought to explore the effects of prolonged antibiotic therapy on microbial stability. To this end stability was compared across three groups: two consecutive samples obtained: 1. during antibiotic treatment or within 30 days after treatment discontinuation (ABX+/ABX+); 2. In the absence of antibiotic treatment for >30 days (ABX-/ABX-) and 3. One during antibiotic treatment and the other in its absence (ABX+/ABX-). Despite a slightly higher (lower Jaccard distance) median stability in ABX-/ABX- compared to the other groups, stability was overall comparable (Fig. 2b). Thus, the intra-personal microbiota stability between time points without antibiotic treatment is just marginally higher compared to stability during antibiotic treatment.

### Microbiome alterations following long-term antibiotic treatment

Our next step was to demonstrate the effect of antibiotic intervention on longitudinal composition of the microbiome and clinical phenotype in patients with pouchitis. Here we noticed that in contrast to IBD [32], but in line with the relatively low degree of intra-personal stability shown in this study, samples from the same patient were not well separated from samples from other patients according to the microbiome composition (Supplementary Fig. 3). Microbial communities from antibiotic-treated samples were strongly separated from non-treated microbiomes (R=0.32, ANOSIM separation, P<0.05, Supplementary Fig. 3), in contrast to the very weak clustering of samples according to clinical phenotype (R=0.06, ANOSIM separation, P<0.05).

One of the most relevant questions during chronic antibiotic therapy is which bacterial species are enriched and which ones are reduced. Thus, we analyzed the species taxonomic profiles of the metagenomes generated with MetaPhlAn2 (Fig. 2c, Supplementary Table 3). *Escherichia coli* (*E. coli*) and two unclassified species of the *Escherichia* and *Pantoea* genera, all belonging to the family *Enterobacteriaceae*, were significantly enriched during antibiotic treatment (Fig. 2c, Mann-Whitney, P<0.05); most notably *E. coli* mean abundance increased from 12.8% (Abx- samples) to 33.8% during antibiotic treatment. *Enterococcus faecium*, an opportunistic human pathogen in patients with severe underlying diseases [33], was significantly increased as well, from 0.7% mean abundance in Abx- to 3.5% in Abx+ samples (Mann-Whitney, P<0.05). Several species from the order *Clostridiales* significantly decreased after chronic antibiotic usage (Mann-Whitney, P<0.05). Most notably, *Faecalibacterium prausnitzii*, a putative anti-inflammatory species [34] which is characteristically reduced in patients with IBD [12,35] as well as in pouchitis [13], was reduced from 3.2% mean abundance to 0.6% in the antibiotic treated samples. *Ruminococcus* spp were also reduced in Abx+ group as previously reported for different species of the *Lachnospiraceae* in pouchitis [13]. Chronic antibiotic usage also resulted in a reduction of several species from *Bifidobacterium* genus, specifically *B. longum and B. animalis*. Interestingly, *Ruminococcus gnavus*, *Bacteroides vulgatus* and *Clostridium perfringens*, all of which were significantly reduced in Abx+ samples (Fig. 2c, Mann-Whitney, P<0.05), had been previously reported to be associated with a higher risk of developing pouchitis after pouch surgery [36]. Overall, our data suggest that long-term antibiotic treatment used to alleviate pouchitis may actually be a double-edged sword. While it diminishes some pro-inflammatory species, it reduces potentially beneficial species, selects for opportunistic facultative anaerobic species and reduces bacterial diversity.

### The majority of commensal bacteria in the microbiome develop fluoroquinolone resistance by point-mutations

Patients with pouchitis in our cohort were treated according to suggested treatment algorithms [6,14] with ciprofloxacin, metronidazole or with both (abbreviated hereafter as C+M). Metronidazole only affects strict anaerobes [37,38], and thus typically shifts the community toward facultative anaerobes, which are inherently resistant to it. In contrast, ciprofloxacin has a broader spectrum of activity [39], and can affect the vast majority of bacteria. C+M was the most common treatment for patients in our study, (20 patients treated with C+M and seven alternated C+M / only ciprofloxacin, Supplementary Table 1c) and should affect all bacteria in the fecal microbiome, unless they have acquired resistance. We therefore focused on metagenomes of the patients treated chronically with C+M and inferred their resistomes.

Known resistance mechanisms to ciprofloxacin (and to other fluoroquinolones [FQ]), primarily involve point mutations in the chromosomal target genes (*gyrA* and *parC*, encoding subunits of gyrase and topoisomerase, respectively), plasmid-encoded drug modifying enzymes (*aac*), and target protection proteins (*qnr*) [40]. We first focused on point mutations, since these are the most common and confer the highest level of FQ resistance [41]. We assembled our metagenomes in order to extract *gyrA* and *parC* alleles and devised a quantitative workflow based on unique multiple sequence alignments for each genus, to infer whether they encode resistant variants (see Methods). Remarkably, most of the dominant genera (*Escherichia*, *Enterococcus*, and *Streptococcus*) in the fecal microbiomes of Abx+ patients had at least one occurrence of a putative resistant allele in either *gyrA* or *parC* (Table 1, Fig. 3a, Supplementary Fig. 4a). For some other genera, resistant alleles were also observed in Abx- patients (Fig. 3b, Supplementary Fig. 4b), indicating that some of these alleles probably do not incur a strong fitness cost and that for these genera (*Bifidobacterium*, *Lactobacillus*, *Dialister*, *Veillonella*, *Bacteroides* and *Prevotella*) background FQ-resistance exists. Genera in which we detected double (in *gyrA*, for a single read) or even triple mutations (*gyrA* + *parC*) appeared almost exclusively in Abx+ microbiomes (Supplementary Table 4). These included *Escherichia*, *Enterococcus*, *Streptococcus*, and *Lactobacillus* (Table 1, Supplementary Table 4). Additionally, we detected FQ-resistant variants of probiotic species (e.g. *Lactobacillus acidophilus, L. delbrueckii, B. longum and B. animalis*) as well as some potential pathogens (e.g. *E. coli*, *K. pneumoniae*, *E. faecium*, *S. gallolyticus, Bacteroides fragilis*) - Table 1. Thus, antibiotics may have unpredictable effects as far as which drug-resistant species they may enrich for.

**Fig. 3.**
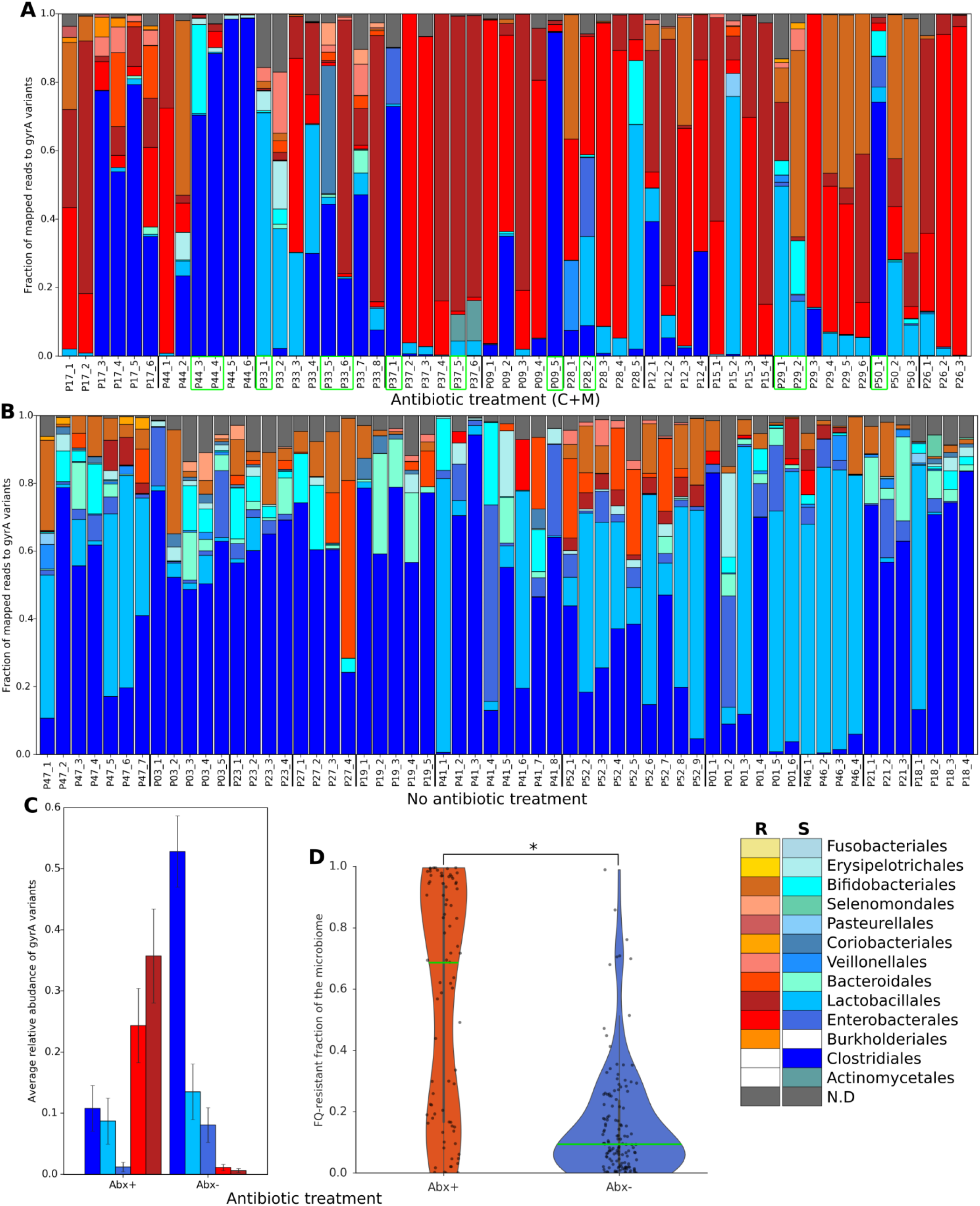
Inferred FQ-resistance of the microbial community over time. The analysis is based on mutation inference of *gyrA* variants for each bacterial genus. The taxa are summarized to order level; “R” signifies all members of that order which carry at least one *gyrA* mutation conferring FQ-resistance (a resistant allele) while “5” signifies that all members carry the sensitive allele (no mutations) and are putatively FQ-sensitive; orders marked in white were below detection. Fecal microbiota from (a) ciprofloxacin and metronidazole antibiotic treatment (C+M) and (b) no antibiotic treatment. Green boxes indicate samples taken during a pause in antibiotic treatment for over 30 days. The plot includes 22 patients with their individual time points (n=115), (c) High abundance orders in the metagenomes according to mutation inference of *gyrA* variants, that significantly differ in abundance between Abx+ and Abx- groups (Mann-Whitney U test, P<0.05, FDR corrected); One sample only from each patient was used in this analysis (n=48) to avoid the potential dependencies between samples and variable number of samples per patient. Error bars indicate the standard error of the mean, (d) FQ-resistant fraction of the microbiome across all samples (assembled metagenomes, n=215) categorized according as either Abx-l- or Abx-, Mann-Whitney U test, *P=1.4e-13. Violin plots whiskers mark observations within 1.5 interquartile range of the upper and lower quartiles.

**Table 1.**
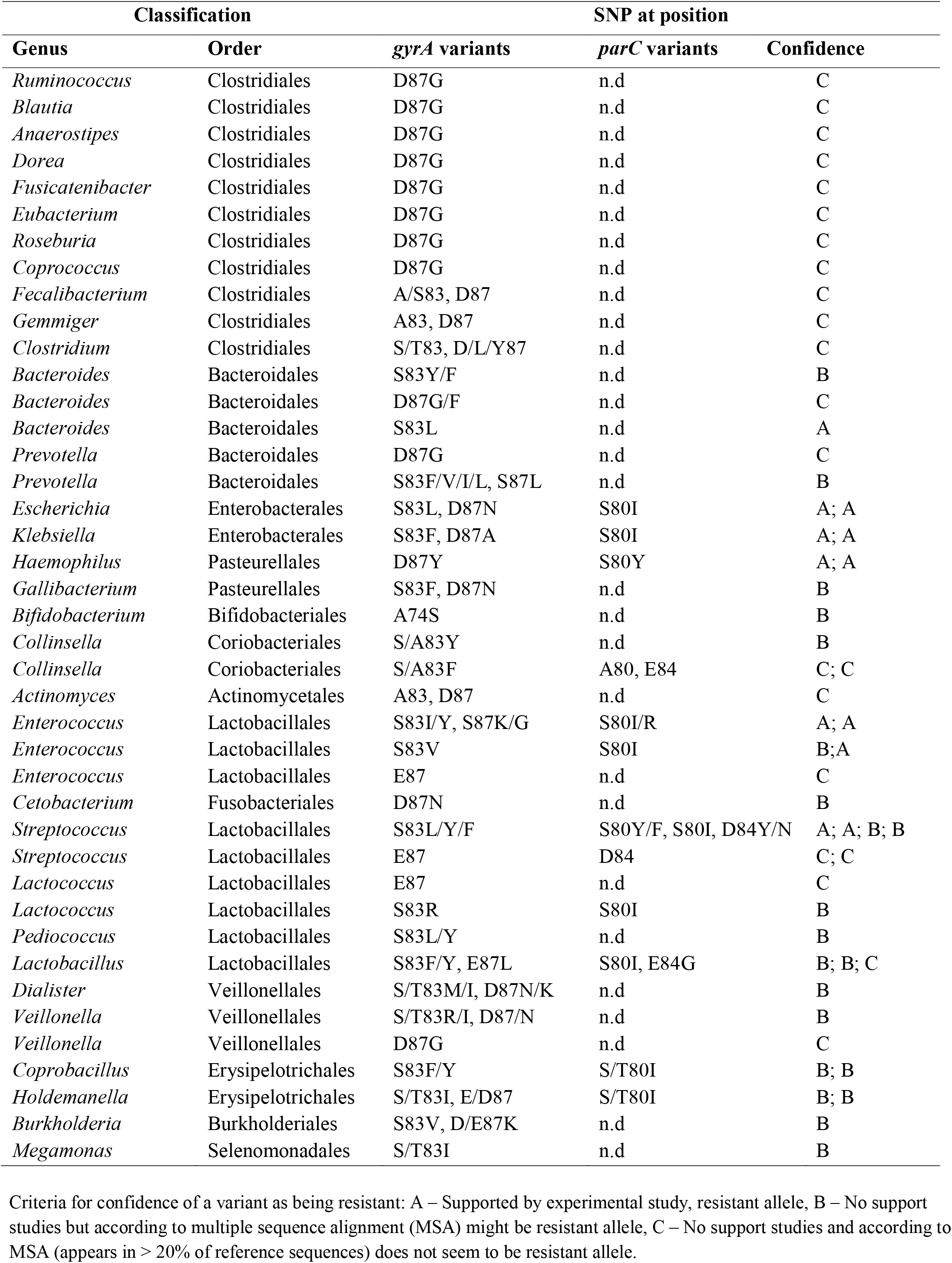
DNA gyrase (*gyrA*) and topoisomerase (*parC*) variants conferring FQ-resistance.

Being able to quantitatively infer putative resistance from metagenomic data enabled us to determine which fraction of the pouch bacterial community and what taxa are FQ-resistant, based on target alleles (Fig. 3, Supplementary Fig. 4). The median fraction of community FQ-resistance in Abx+ samples was 72%, compared to 9.4% in samples obtained in Abx- conditions (Mann-Whitney, P=1.4×10^-13^, Fig. 3d, Supplementary Table 5). Furthermore, in Abx+ microbiomes the mean abundance of resistant *Enterobacterales* and *Lactobacillales* was 24.4% and 35.7%, respectively, in contrast to only 1.13% and 0.6% in Abx- microbiomes (Mann-Whitney, P=1.5×10^-5^ for *Enterobacterales*, P=8.4×10^-6^ for *Lactobacillales*, Fig. 3c, Supplementary Table 5). Lastly, FQ-sensitive *Clostridiales* had mean abundance of 52.8% in non-treated samples, but decreased to 10.8% during C+M treatment, probably due to the strong effect of the combination of ciprofloxacin with metronidazole that is effective against obligate anaerobes (Mann-Whitney, P=1.7×10^-5^, Fig. 3c, Supplementary Table 5). This implies that continuous antibiotic treatment with C+M generates a microbiome dominated by multiple FQ-resistant facultative anaerobes.

### Prolonged treatment with concomitant ciprofloxacin and metronidazole enriches the microbiome for mobile resistance genes

FQ-resistance may also be conferred by mobile resistance genes, and can be encoded on the mobile elements that carry resistance genes to additional antibiotics. This may have clinical implications for the patients and those that come in contact with them that are broader than resistance to the administered antibiotics. To obtain an overall bacterial resistome in our samples, we profiled the assembled metagenomes for all antibiotic resistance genes (ARG) based on the Comprehensive Antibiotic Resistance Database (CARD) for antibiotic resistance ontology (see Methods, Supplementary Fig. 5, Supplementary Table 6). Specific aminoglycoside acetyl transferases (*aac*) families can inactivate multiple antibiotics, including FQ, and provide low level resistance to ciprofloxacin [40]. We detected *aac(6’)-Ib-cr* in 37 samples from 13 patients (Table 2), of which only a single patient (one sample) was not treated with ciprofloxacin in the year prior to sampling (Fisher exact test, P=2.5×10^-10^). *aac(6’)-Ib-cr*, along with genes conferring beta-lactam and chloramphenicol resistance (*oxa-31* and *catB3*, respectively), are often located on integrons that are encoded by plasmids of *Enterobacterales*, such as *E. coli* and *Klebsiella pneumoniae* [42]. Accordingly, in our assembled contigs *aac(6’)-Ib-cr* was always co-localized with *oxa-31* and *catB3*, as well as with additional genes conferring resistance to trimethoprim and rifamycin (Table 2).

**Table 2.** Mobile co-localized ARG on assembled contigs from the metagenomes conferring multiple resistance.

| Assembled contigs |  |  |  | Best BLASTN Match |  |  | Prevalence |  |
| --- | --- | --- | --- | --- | --- | --- | --- | --- |
| Name | length (Kb) | ARG | Drug class | Annotation | GenBank # | Identity (%) | # of patients | # of samples |
| NODE_7209 | 4.9 | *qnrS1*, *TEM-215* | FQ, BL | *K. pneumoniae* plasmid pK245, *E. coli* plasmid pE66An | DQ449578.1, HF545433.1 | 99, 99 | 1 | 3 |
| NODE_15824 | 2.2 | *qnrS1* | FQ | *K. pneumoniae* plasmid pSg1-NDM, *E. coli* plasmid pE66An | CP011839.1, HF545433.1 | 99, 99 | 1 | 1 |
| NODE_4266 | 8.75 | *qnrS1*, *TEM-215* | FQ, BL | *K. pneumoniae* plasmid pKPC-CR-HvKP267, *E. coli* plasmid pE66An | MG053313.1, HF545433.1 | 99, 99 | 1 | 2 |
| NODE_325 | 46.6 | *qnrS1*, *TEM-1* | FQ, BL | *E. coli* plasmid pEQ2, *Shigella flexneri* 1a strain 0228 plasmid | KF362122.2, CP012734.1 | 99,99 | 1 | 1 |
| NODE_6846 | 3.3 | *qnrS2* | FQ | *K. pneumoniae* plasmid pKPSH231 | KT896503.1 | 100 | 1 | 2 |
| NODE_884 | 19.1 | *qnrS1*, *TEM-1*, *TEM-215*, *sul2*, *APH(3'')-Ib*, *APH(6)-Id* | FQ, BL, SA, AM | *E. cloacae* plasmid pEC27-2 | CP020091.1 | 99 | 1 | 1 |
| NODE_671 | 23.6 | *qnrB4*, *DHA-1*, *sul1* | FQ, BL, SA | *K. pneumoniae* plasmid pHM881QN | LC055503.1 | 99 | 1 | 5 |
| NODE_37316 | 1.6 | *qnrB4* | FQ | *E. coli* strain plasmid A, *Cronobacter sakazakii* plasmid p505108-MDR | CP010239.1, KY978628.1 | 99, 99 | 1 | 2 |
| NODE_1711 | 11.7 | *qnrS1*, *TEM-1* | FQ, BL | *E. coli* plasmid pEQ2, *Shigella flexneri* 1a strain 0228 plasmid | KF362122.2, CP012734.1 | 100, 100 | 3 | 4 |
| Total |  |  |  |  |  |  | 11 | 21 |
| NODE_12591 | 3 | *AAC(6')-Ib-cr*, *OXA-31*, *catB3* | FQ, BL, CP | *E. coli* plasmid *pHS13-1-IncHI2*, *E. cloacae* plasmid pMRVIM0813 | CP026492.1, KP975077.1 | 100, 99 | 4 | 14 |
| NODE_7063 | 2.4 | *AAC(6')-Ib-cr*, *OXA-31*, *cat* | FQ, BL, CP | *E. coli* plasmid unitig_3j3rc_linear, *K. pneumoniae* plasmid INF274 | CP021937.1, CP024573.1 | 100, 100 | 7 | 11 |
| NODE_2765 | 5.25 | *AAC(6')-Ib-cr*, *OXA-31*, *catB3*, *arr-3* | FQ, BL, CP, rifamycin | *E. coli* plasmid *pHS13-1-IncHI2*, *E. cloacae* plasmid pMRVIM0813 | CP026492.1, KP975077.1 | 100, 99 | 1 | 6 |
| NODE_2476 | 9.2 | *AAC(6')-Ib-cr*, *aadA*, *aadA25*, *cmlA6*, *dfrA12* | FQ, AM, ME, trimethoprim | *E. coli* plasmid p19M12 | KY689632.1 | 99 | 1 | 4 |
| Total |  |  |  |  |  |  | 13 | 35 |

FQ – fluoroquinolone, BL – beta-lactam, SA – sulfonamide, AM – aminoglycoside, CP – chloramphenicol, ME – multidrug efflux.

*qnr* (quinolone-resistance) genes encode target-protection proteins that hinder the binding of ciprofloxacin to topoisomerases, conferring moderate resistance to FQ and are usually plasmid encoded. We detected *qnrS1* in 25 samples from 18 patients and *qnrB4* in 18 samples from 12 patients (Table 2). Surprisingly, these *qnr* genes were not significantly enriched in Abx+ samples. Similar to *aac(6’)-Ib-cr*, the contigs on which these *qnr* genes occurred matched sequences from known plasmids of *E. coli* and *K. pneumoniae* (Table 2), and also encoded beta lactamases belonging to the TEM family [43].

Healthy human gut microbiomes may contain antibiotic resistance genes even in untreated individuals [21,22]. Indeed, many such genes in our dataset were not treatment-associated. Beta-lactam, polymyxin, macrolide, and tetracycline resistance genes, as well as efflux pump genes were commonly encountered in metagenomes of both Abx+ and Abx- samples (Supplementary Fig. 5). Vancomycin and aminocoumarin resistance genes were decreased in Abx+ samples, probably reflecting the decrease in species from *Clostridiales* order that harbour these genes (Supplementary Table 6, Mann-Whitney, P<0.05). In contrast, prolonged antibiotic treatment also enriched the microbiome in trimethoprim resistance genes (*drfA, drfE* and *dfrG*) (Mann-Whitney, P<0.05, Supplementary Fig. 5, Supplementary Table 6), which showed 100% sequence similarity to *E. coli* and *E. faecium*, and are probably increased due to their co-localization with *aac*-encoding plasmids (see above). Importantly, we detected extended-spectrum β-lactamases genes encoding for CTX-M-14/15 [44] (100% protein identity to *E. coli*) in 31 samples from 17 patients (Supplementary Table 6c), only six of which were taken from patients who were not treated with C+M during sample collection or in the year prior to sampling (Fisher exact test, P=0.0005). Moreover, we found that there are significantly more positive correlations between dominant species abundance and resistance genes in Abx+ (285) compared to Abx- (185) samples (Fisher exact test, P=5.9×10^-8^, Supplementary Fig. 6), which suggests that there is a general enrichment of resistance genes in the microbiome following specific antibiotics treatment (of C+M). To infer how long does antibiotic treatment-associated enrichment of resistance genes persist, we divided the samples based on the time elapsed from last antibiotic treatment. In the Abx+ samples, the fraction of resistance genes in the microbiomes was 30% higher (Mann-Whitney, P<0.05, Supplementary Fig. 7a) than in Abx- (0.71% from the total reads mapped to assembled resistance genes in Abx+ compared to 0.55% in Abx-). In contrast, no difference was observed in the fraction of resistance genes in samples that were antibiotic-free for 1-6 months compared to those obtained when patients were free of antibiotics for over six months (Supplementary Fig. 7b). Thus, most antibiotic resistance genes enrichment diminishes gradually in the months following treatment discontinuation.

Taken together, our results suggest that although most FQ-resistance is chromosomal, chronic treatment with a combination of C+M is associated with a higher risk of carriage of multidrug resistance plasmids and a general enrichment with antibiotic resistance genes. Thus, treating with this drug combination may inadvertently increase collateral resistance to non-related antibiotics for up to several months after antibiotics discontinuation.

### Bacterial density is moderately decreased following prolonged antibiotic treatment in patients with pouchitis

Antibiotics are the mainstay therapy for pouchitis, which is considered to be the most antibiotic-responsive IBD. Yet the mechanisms underlying clinical response are unclear [6,14,45]. One possible explanation for the clinical efficacy could be that by using broad spectrum antibiotics (e.g. C+M), the numbers of bacteria in the pouch are greatly reduced, resulting in lower exposure to bacteria-derived antigens ("reducing antigenic load"), thereby dampening inflammation. An alternative explanation is that there are *specific* bacterial taxa that greatly contribute to inflammation ("pathobionts") and that these taxa are suppressed by antibiotics. To distinguish between these scenarios would require quantification of bacterial concentration in the pouch, reflected by the fecal concentration. Currently, fecal bacterial concentration (bacterial density) is measured by qPCR [28] or flow cytometry [46]. We used shotgun metagenomics, which does not require complicated cell labeling, extensive and lengthy calibration of qPCR-based measurement and primer biases, as a proxy to bacterial concentration. For that end we developed an index based on the ratio of bacterial to non-bacterial (derived from human host or from viruses) reads from the metagenomes [the B/(H+V) ratio, see Methods, Supplementary Table 2a]. Since the levels of human and viral DNA in faeces should be relatively independent of those of bacterial DNA, and shotgun metagenomics sequences all DNA irrespective of its origin, a higher fraction of reads derived from bacteria should be strongly correlated with bacterial concentration in the sample. Human reads fraction ranged widely from 0.18% to 90.5%, with a mean of 8.3% and virus reads ranged from 0.07% to 27.7%, with a mean of 1.3% of the total reads in the metagenomes (Supplementary Table 2a). To test the agreement between qPCR measurements and our metagenome-inferred bacterial density, we plotted B/(H+V) vs. qPCR-based bacterial density for 11 samples taken from patients with a pouch (Fig. 4a). A strong linear correlation (Pearson r=0.84, P=0.0012) between qPCR and B/(H+V) ratios was observed, indicating that our metagenome-derived ratio well represents the bacterial density in fecal samples of patients with a pouch. We then compared bacterial density computed by B/(H+V) across our samples, grouped according to antibiotics treatment and clinical phenotype of the pouch. Samples obtained during antibiotic therapy had a median B/(H+V) ratio 1.9 times lower compared to Abx- samples (Fig. 4b). This moderate reduction is comparable to previous culture-based findings observed in patients with pouchitis treated with a different antibiotics combination for 15 days [47], as well the reduction observed using qPCR in mice treated with C+M [28]. Moreover, microbiomes with higher species diversity were associated with higher bacterial density (Fig. 4c, Spearman r=0.28, P=2.6×10^-5^), as observed previously in humans with an intact colon using flow cytometry [46]. Lastly, we observed that patients with pouchitis (Chronic, CLDP and some with recurrent acute pouchitis) had lower bacterial density compared to patients with a normal pouch and FAP (Fig. 4d). This may reflect their antibiotic usage but also pouch inflammation, both of which can decrease the bacterial density in the pouch.

**Fig. 4.**
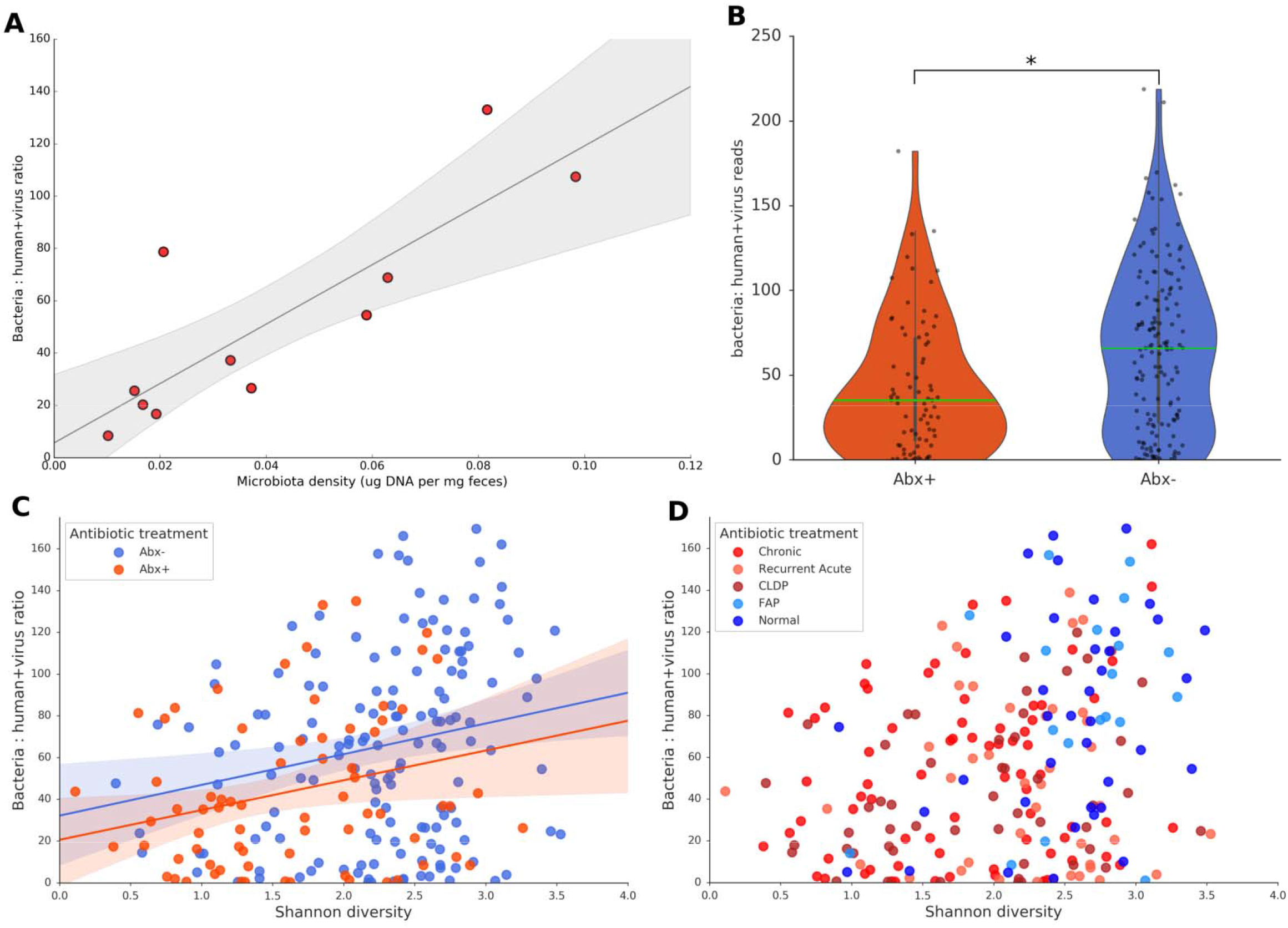
Metagenome-inferred bacterial density approximated by the ratio of bacterial to human+viral reads [B/(H+V)]. (a) High correlation between microbiota density (qPCR) and bacteria : human-virus reads ([B/(H+V)]_r_ metagenomics) for 11 samples for which both data is available (Pearson r=0.84, P=0.0012 / Spearman r=0.85, P=0.0008). (b) Bacterial density is moderately but significantly reduced following long term antibiotic treatment, based on B/(H+V) ratio (see Methods); Mann-Whitney test P=7.5e-4 (c) Association between bacterial density estimated by [B/(H+V)] and species diversity in the gut microbiota across all samples in patients with pouchitis; Spearman r=0.28, P=2.6e-05. Samples are colored as either Abx+ or Abx-. (d) The same analysis as in (c) but samples are colored according to the clinical phenotype of the pouch. For (b-d) all the samples were analyzed (n=234). Line represents least squared error (linear) regression fit with a 95% confidence interval (20000 bootstrap resamples). Violin plots whiskers mark observations within 1.5 interquartile range of the upper and lower quartiles.

Previous work argued that antibiotic treatment failure in patients with pouchitis was due to ciprofloxacin-resistant *Enterobacteriaceae*, which were present in all those patients [48]. Here we show that C+M treatment indeed powerfully selects for such FQ-resistant bacteria that were present in varying abundances in nearly all samples of C+M-treated (65 out of 71, Supplementary Table 5) patients, however, all these patients maintained clinical remission at time of treatment. Thus, the development of FQ-resistant bacteria is not the major explanation for failure of antibiotic treatment. To expand on this, we surveyed our metagenomes for *Enterobacteriaceae-*derived enterotoxin genes (SPATE – extracellular proteases secreted by *Enterobacteriaceae*, see Methods, [49]) and observed that only an average of 3.2% of the samples had putative enterotoxins, in contrast to toxins important in extra-intestinal infections that were common (25%) in the patient metagenomes (Fisher exact test, P=1.3×10^-37^, Supplementary Table 7). We also compared the prevalence of enterotoxins in Abx+ vs. Abx- samples and found no significant difference. Thus, prolonged antibiotic treatment does not select for enterotoxigenic *Enterobacteriaceae*, an additional support for the success of the treatment. Therefore, antibiotic resistance of “usual suspects”, such as *Enterobacteriaceae*, does not predict treatment failure.

## CONCLUSION

Antibiotic resistance is an emerging global health problem [50], and there is well-justified pressure to minimize the spread of resistant bacterial species by better antibiotic stewardship. Long-term antibiotic therapy can place the patient and her or his environment under constant risk of transmitting resistant potentially-pathogenic strains. Here we took advantage of a unique cohort of patients that have detailed longitudinal follow up to examine resistome dynamics, following such long-term treatment. We show that long-term antibiotic treatment results in a relatively stable microbial community that is reduced only two-fold in its bacterial density. While this may seem surprising, the pervasive antibiotic-resistant alleles that we observed in those microbiomes can explain these observations. We also showed collateral resistance, which can be attributed to co-localization of different resistance genes on the same mobile element. This collateral resistance represents a concern for patients and the community due to potential horizontal gene transfer of such elements to other bacteria or the transmission of the resistant strains themselves.

Collectively, our data show that prolonged antibiotic treatment modestly reduces the bacterial density and is unlikely to reduce the overall immune exposure to bacteria-derived antigens. This contradicts the accepted assumption that the rationale for treating IBD, specifically pouchitis, with antibiotics is reduction of antigenic load. We therefore suggest that suppression of pro-inflammatory pathobionts is a more likely explanation for antibiotic treatment success in pouchitis and that different patients may have different pathobionts. This may parallel the success of some antibiotic combinations [51] in the treatment of active UC [52]. We now suggest that clinical management of pouchitis should therefore focus on identifying and eliminating these pathobionts via targeted eradication regimes, such as those used for *Helicobacter pylori* [53]. Such treatments will improve upon the current standard of care that involves chronic treatment that inevitably leads to collateral antibiotic resistance.

## MATERIALS AND METHODS

### Study design and cohort details

Patients after pouch surgery were recruited at the Comprehensive Pouch Clinic where an IBD-oriented gastroenterologist and a colorectal surgeon routinely followed up all patients. Pouch disease behaviour (phenotype) was defined as normal, acute/recurrent-acute, chronic or Crohn’s-like disease of the pouch (CLDP) for patients who underwent surgery due to UC, as previously defined [4]. Patients undergoing pouch surgery due to familial adenomatous polyposis (FAP) were recruited as well. Briefly, a normal pouch was defined as no pouchitis during the past 2 years and no antibiotic therapy. Acute/recurrent-acute pouchitis was defined as a flare of pouchitis responding to a short (usually 2 weeks) antibiotic therapy, or up to 4 flares/year, respectively. Chronic pouchitis was defined as >4 pouchitis flares/year or the need for chronic administration of antibiotic or IBD specific anti-inflammatory therapy for more than 1 month. CLDP was defined as having pouch-perianal disease, pouch strictures, or long segments of proximal small intestinal inflammation.

Demographics and clinical data were collected during clinic visits. These included type of antibiotics used, treatments duration, fecal calprotectin and pouch disease phenotype (Supplementary Table 1a,b). Fecal samples were collected in sterile cups during each visit and immediately frozen at −80 °C until processing. Stool samples were collected longitudinally over the course of up to 5.9 years (intervals ranging from one month to one year between samples). In total, 49 patients after pouch surgery were included in this study and their 234 corresponding samples were used in downstream analysis (after quality control). The patients were treated with several antibiotics regimes as follows: 20 patients treated with a combination of C+M antibiotics, seven alternated between C+M or only ciprofloxacin, five were treated with ciprofloxacin, one was treated with metronidazole and 16 were not treated with antibiotics during follow-up time (Supplementary Table 1c,d). For downstream statistical analysis samples were grouped in two ways: First, Abx+, samples obtained during antibiotic treatment or up to one month post treatment; Abx-, samples taken from patients not treated with antibiotics for at least one month. Second, according to the time that elapsed since the last antibiotic use as follows: 1 - obtained during ongoing treatment, 2 – obtained after treatment was stopped but less than a month post treatment, 3 – obtained more than a month post treatment but less than half a year after treatment was stopped, 4 – obtained over half a year or longer post treatment.

### Genomic DNA extraction and shotgun metagenomic sequencing

Fecal samples were thawed at room temperature and total genomic DNA was extracted using the PowerLyzer PowerSoil DNA Isolation Kit (MOBIO, Carlsbad, CA) using the kit extraction protocol. The OMNI Bead Ruptor 24 Homogenizer (OMNI International, Kennesaw, GA) was used for sample homogenization at the following settings: speed 5.65 m/s, cycles 02, run time 0:45 minutes, and dwell time 0:30 minutes. Extracted DNA samples were stored at −80°C.

Genomic libraries were prepared with the Nextera XT library preparation kit using approximately 1ng total DNA per sample (DNA concentrations verified by Qubit fluorometry). Metagenomic sequencing was done using Illumina NextSeq500 paired-end (2 x 150 bp reads) at DNA Services Facility, University of Illinois, Chicago, IL, USA; 234 samples were obtained (after removal of low sequencing depth samples). The sequencing depth (mean±sd) was 1.02±0.24 Gbp after quality control per metagenomic sample (Supplementary Table 2a). Quality control step was done with Trimmomatic v0.36 [54] using default parameters, which consisted of low-quality reads filtering and removal of low score bases and short reads.

### Taxonomic profiling of the metagenomes

Taxonomic profiling was performed using MetaPhlAn2 classifier v2.6.0 [55], which classified metagenomic reads by mapping to a database of clade-specific marker genes. MetaPhlAn2 was run on all the samples that passed quality control and had sufficient number of bacterial reads (n=225) with the following parameters changed from default: *–tax_lev* set to ‘s’ (classify taxonomy to species level), *–ignore_virus* and *–ignore_eukaryotes* to ignore viral and eukaryotes reads respectively. The output species relative abundance tables from each sample were merged together. The following downstream analysis used this merged species table: Shannon diversity, microbiota stability analysis, Principal coordinate analysis, differential enrichment analysis (LefSe v1 [56]), and Pearson correlation between taxa and ARGs. Briefly, Shannon diversity and Chao richness were calculated using *diversity* and *specnumber* functions in R package vegan. Microbiota stability was calculated with Jaccard distance metric using *cdist* function in Python package SciPy, the input table was first transformed to binary (presence/absence) before applying the function. Principal coordinate analysis was performed with either Bray-Curtis or Jaccard distance matrices using *cmdscale* function in R. To detect which bacterial species were differentially enriched in the grouped samples, LefSe was run with default parameters on the merged species table. Pearson correlation coefficients between 25 abundant bacterial taxa and 162 ARGs (see below) across all the metagenome assembled samples (n=215, see below) were calculated using *corrcoef* function in Python package NumPy.

### Metagenome assembly and antibiotic resistance genes quantification

*De novo* metagenome assemblies were done with SPAdes assembler v3.11.0 [57]; all the follow-up metagenomes belonging to individual patients were pooled together to increase sequenced reads coverage and assembled together. SPAdes was run in paired-end mode with the following parameters: *–careful* (to minimize the number of mismatches in the contigs), *-k* 21, 33, 55, 77, 99, 127 (k-mer lengths, SPAdes’ use of iterative k-mer lengths allows benefiting from the full potential of long paired-end reads). A summary of all metagenome assemblies’ statistics is found in Supplementary Table 2b. Assembled contigs smaller then 1kb were discarded. The rest of the contigs were used for open reading frame prediction using the gene finding algorithm Prodigal v2.6.3 [58]. Annotation of the predicted genes was done using two approaches; ARGs were identified with Resistance Gene Identifier (RGI) v3.1.1 [59,60] and all prokaryotic genes in the assemblies were annotated using Prokka v1.12 [61]. Briefly, RGI uses the predicted protein-coding genes and annotate them based on BLASTP searches (cutoff of e^-30^) against the curated protein sequences in Comprehensive Antibiotic Resistance Database (CARD [59] download April 2017). For this analysis of resistance genes we excluded protein variant models (SNPs) results and kept only protein homolog models. Prokka uses BLASTP searches against UniProtKB followed by more sensitive hmmscan (HMMER3) scan against hidden Markov model databases to assign function to protein-coding genes and was run with *–metagenome* parameter. To obtain sequence taxonomy, the identified ARGs were used as a query for BLASTP search against NCBI reference proteins (RefSeq) database (download 15 May 2017).

In order to quantify the ARGs (and all prokaryotic genes), the metagenomic reads of each sample were separately mapped to the assembled contigs using Bowtie2 short reads mapper v2.2.9 [62] in *–very-sensitive-local* mapping mode and SAMtools v1.3.3 [63]. ARGs (and all prokaryotic genes) coverage (counts of mapped reads) was calculated using BEDtools v2.26.0 [64] function *multicov* provided with a mapping file (.bam) and gene coordinates file (.gff) for each sample. For downstream statistical analysis, ARGs from all the samples were merged together, rare resistance genes (present in <10% of the metagenomes) were filtered out and the rest were normalized with *cumNorm* function (cumulative sum scaling normalization [65]) from R package *metagenomeSeq*. In total 272 unique ARGs were identified and after rare genes filtration 162 ARG left.

In addition, in order to detect presence of genes encoding for potential toxins, the metagenomic reads were mapped against a reference genes set of SPATE (Serine Protease Autotransporters of *Enterobacteriaceae* [49]). Briefly, we used five enterotoxins (*eatA*, *epeA*, *espC*, *espP*, *pet*) and five extra-intestinal toxins (*hbp*, *pic*, *sat*, *tsh*, *vat*) as a reference (for genes description and mapping summary see Supplementary Table 7). The mapping and quantification was done with Bowtie2 and SAMtools as mentioned previously. Finally, the gene counts were transformed to presence / absence to perform Fisher exact test.

### Single nucleotide polymorphism analysis for fluoroquinolone target genes to infer resistance for commensal bacteria

To be able to infer resistance to FQ antibiotics in the microbiome, we devised the following workflow: From the metagenome assemblies we extracted all the DNA gyrase subunit A, B (*gyrA*, *gyrB*) and topoisomerase IV subunit A, B (*parC*, *parE*) which are the FQ target genes. BLASTP search against NCBI RefSeq or NR databases was used to assign taxonomic annotation to the genes. For each bacterial genus *gyr* and *par* identified, we built a separate multiple sequence alignment of the amino-acid *gyr*/*par* sequences of reference taxa downloaded from NCBI protein and compared it to *gyr*/*par* genes from our assemblies. The quinolone resistance determining region (where mutations can arise, resulting in amino-acid substitutions which alter the target proteins leading to drug resistance) was examined to identify amino-acid substitutions which can differ between taxa. Each potential gene variant was scored as follows: A – supported by an experimental study with isolated strains, a resistant allele; B – no support studies but appears in < 20% of the reference sequences in the alignment, potential resistant allele; C - no support studies and is common in the reference alignment (>=20%), does not seem to be resistant allele. For the FQ-resistance analysis done in this study, only variants scored as A and B were considered as FQ-resistant. Although all the four target genes were examined (*gyrA*, *parC*, *gyrB*, *parE*), we found a high number of potential resistance variants only in *gyrA* and *parC*, which are considered as the primary target genes for fluoroquinolones for most bacterial species. All the identified *gyrA* and *parC* variants in our metagenome assemblies appear in Table 1. Abundance information for *gyrA* / *parC* was obtained using Bowtie2 as mentioned in the previous section. In addition, we wished to validate that in cases where two mutations were detected in a specific *gyrA* sequence, it originated from a single strain. FQ-sensitive *E. coli gyrA* nucleotide sequence was used as a reference and the metagenomic reads of each sample were separately mapped against it with Bowtie2. Detection of variants between the reads and the reference sequence (variant calling) was done using SAMtools *mpileup* in conjunction with BCFtools *call* v1.7-1 [66]. The double mutations in *E. coli gyrA* were found to cover single reads (in samples where *E. coli* was abundant enough to do variant calling), confirming that the double mutant *gyrA* variant is a single strain.

### Bacterial density analysis through metagenomics

We have developed a method to measure relative bacterial density directly from metagenomic samples by measuring the ratio of bacterial to non-bacterial (human host and virus) reads, termed hereafter B/(H+V) ratio. To quantify human reads in the metagenomes, we used GRCh38 human genome (download from NCBI Human Genome Resources) as a reference and mapped the metagenomic reads against it using Bowtie2 and the number of mapped reads was counted. To quantify virus reads, all available viral genomes from NCBI viral genomes database (9556 genomes, downloaded November 2017) were used as reference and quantified as mentioned above. Both human and virus reads were filtered by mapping quality parameter (MAPQ field in .bam alignment file) and only reads with MAPQ >=10 (high probability of accurate mapping) were retained prior to quantification. A mean of 15.4% of human and 35.6% of virus reads were discarded after filtration. Bacterial reads were quantified as follows; the metagenomic reads were used as a query for translated nucleotide BLASTX search against NCBI NR database (download June 2018) using DIAMOND [67] with the following parameters changed from default: *–evalue* set to 0.1, *–max-target-seqs* set to 1 (one hit per read) and *–taxonmap* added. The latter parameter was used to map NCBI protein accession numbers to taxonomy ID (tax_id) from the NCBI Taxonomy database in order to obtain taxonomy information for each read. The taxonomy mapping file can be downloaded from: ftp://ftp.ncbi.nlm.nih.gov/pub/taxonomy/accession2taxid/prot.accession2taxid.gz. Thus for each metagenomic read, superkingdom level was extracted and only ‘Bacteria’ classified reads were counted. Finally, to obtain B/(H+V) ratios for each sample, we divided the number of bacterial reads by the sum of human and virus reads. For the B(H+V) quantification of the metagenomes see Supplementary Table 2a.

### Statistical analysis

Mann-Whitney rank tests, Kruskal-Wallis H-tests, Fisher exact tests, Spearman rank-order correlations and Pearson correlations were performed using *mannwhitneyu*, *kruskal*, *fisher_exact, spearmanr and pearsonr* functions, respectively in Python package SciPy. All reported P-values were adjusted for multiple hypotheses testing using false discovery rate (Benjamini/Hochberg method), with *fdrcorrection0* function in Python package Statsmodels. Multiple comparisons corrections were done after Kruskal-Wallis tests using Dunn’s test (multiple pairwise comparisons among samples groups), with *dunn.test* function in R package dunn.test. Analysis of similarities (ANOSIM) test was done with *anosim* function in R package vegan.

To model the effects of antibiotics and of clinical data on species richness in the microbiome, a generalized linear mixed model (GLMM) was used with Poisson distribution fit by maximum likelihood estimation and log link function. The following predictors were used: antibiotic treatment category (time elapsed since treatment, 1-4), accumulated antibiotic days, clinical pouch phenotype (1 – chronic/CLDP, 2 – recurrent acute, 3 – normal/FAP), calprotectin, patient age, ever VSL-user (yes/no) and ever anti-TNF treatment (yes/no). All predictors were centered and scaled to account for the different variables scales using the function *scale* in base R. Individual patient was specified as random effect. The model was implemented using *glmer* function in R package lme4.

### Data availability

#### Code availability

All the plots, statistical methods and custom scripts in this article were created using open-source software Python, R and Linux Bash. Python software is available at (www.python.org/downloads) and packages are maintained on PyPi repository (https://pypi.org). R software and packages are available on CRAN repository (https://cran.r-project.org).

## SUPPLEMENTARY MATERIALS

Fig. S1. Shannon diversity of the microbiota in patients with a pouch.

Fig. S2. No effect of cumulative antibiotics usage on microbiome diversity.

Fig. S3. Principal coordinate analysis (PCoA) of the bacterial taxonomic profile of the metagenomes at the species level.

Fig. S4. FQ-resistome dynamics of the microbial community over time.

Fig. S5. Antibiotic resistance genes (ARG) profile across all assembled metagenomic samples, collapsed into drug classes according to CARD antibiotic resistance ontology.

Fig. S6. Pearson correlation between 25 most abundant bacterial species profiled by MetaPhlan2 classifier and the assembled ARG in the microbiomes.

Fig. S7. Fraction of ARG in the microbiome is enriched during antibiotic treatment.

Table S1. Study cohort demographics and samples metadata.

Table S2. Bacteria-Human-Virus ratios and metagenome assembly stats.

Table S3. Taxonomic profiles MetaPhlAn2.

Table S4. Species with 2-3 *gyrA*-*parC* mutations.

Table S5. FQ-Resistance Sensitive microbiome.

Table S6. Antibiotic resistance genes profiles.

Table S7. SPATE genes mapping.

## ACKNOWLEDGMENTS

The authors thank Jeremiah Faith of Mount Sinai, New York for providing additional data; Stefan Green of the University of Illinois at Chicago, for his continued expert help in the metagenomic sequencing; Karin Yadgar and Keren Zonensain of Rabin Medical Center, Tel-Aviv for help with obtaining clinical data.

## AUTHORS CONTRIBUTIONS

V.D., U.G. and I.D. conceived and designed the study; V.D. developed the bioinformatic analysis pipelines and analyzed the data; L.R. analyzed data; N.B., K.R., L.G., and I.D. collected and analyzed clinical data; H.T. and I.D. enrolled and examined the patients. V.D., L.R., U.G. and I.D. wrote the paper and all authors read, discussed, and approved the final manuscript.

